# Modulating Neural Tracking of Speech in Infants

**DOI:** 10.64898/2026.07.01.735811

**Authors:** Laura Fernández-Merino, Mikel Lizarazu, Nicola Molinaro, Marina Kalashnikova

## Abstract

Neural oscillations synchronize to the rhythmic structure of speech, a process known as cortical tracking that is thought to support speech perception and language acquisition. Individual differences in cortical tracking during infancy predict later language outcomes, yet little is known about whether this neural mechanism can be shaped by recent auditory experience early in development. Here, we tested whether brief exposure to structured musical rhythms modulates cortical tracking of speech in Spanish–Basque bilingual infants at 6 and 10 months of age. Infants listened to speech preceded by either temporally regular musical sequences that mirrored the rhythmic structure of the speech signal or rhythmically irregular musical sequences. Cortical tracking was measured using electroencephalography and quantified as speech–brain coherence in the delta and theta frequency bands. Regular musical sequences enhanced subsequent cortical tracking of speech, but the effects varied across age, language, and frequency band. In Spanish, rhythmic priming enhanced delta-band tracking at 10 months of age, whereas in Basque it enhanced theta-band tracking at both 6 and 10 months. These language-specific effects suggest that rhythmic priming interacts with the temporal properties of individual languages and infants’ developing linguistic experience. Together, the findings demonstrate that cortical tracking is a highly flexible neural mechanism during infancy and can be rapidly modulated by structured rhythmic input. Thus, we identify rhythmic experience as a potential pathway through which infants’ auditory environment shapes neural speech processing during language development.

## 1. Introduction

Speech is characterized by quasi rhythmically structured patterns that provide essential cues for segmentation and comprehension. Neural oscillations naturally align with these rhythmic structures, a process known as cortical tracking of speech. This process reflects the adaptation of the brain oscillations to the temporal regularities of the external acoustic signal (see Meyer, 2018; and Molinaro, 2025), corresponding to different levels of the linguistic hierarchy, including phrases and words, syllables, and individual phonemes. Studies with adult participants have shown that cortical tracking supports the segmentation and identification of these linguistic units in speech (Meyer, 2018). Critically, there is emerging evidence that cortical tracking can be successfully modulated by immediate brief exposure to musical rhythms (Fernández-Merino et al., 2025; Falk et al., 2017), with the potential to enhance speech processing. Investigating whether such modulation can also be achieved in infancy is particularly important because the first year of life represents a critical period for the development of speech segmentation and the neural encoding of linguistic structure. Here, we assess this possibility by directly testing whether rhythmic exposure can enhance cortical tracking of speech during infants’ first year of life.

Cortical tracking of the speech envelope, which carries crucial acoustic information required for perceptual encoding and decoding at multiple temporal scales, has been detected already at birth, for both infants’ native and non-native languages (Ortiz-Barajas et al., 2023). This suggests that prenatal exposure to rhythmic cues and envelope features, which are already perceived in utero (Gorina-Careta et al., 2024), lays the groundwork for this mechanism. Over their first months, infants become increasingly attuned to the rhythmic properties of their native language, a process reflected in developmental changes in cortical tracking responses (e.g., Attaheri et al., 2022; Kalashnikova et al., under review). Importantly, different linguistic timescales have been associated with distinct oscillatory frequencies, with slower delta-band activity (∼1-3 Hz) often linked to the tracking of stressed syllables and prosodic structure, and faster theta-band activity (∼4-8 Hz) associated with syllabic processing. In a longitudinal study with English-learning infants from 4 to 11 months, Attaheri et al. (2022) showed stronger delta-band tracking of nursery rhymes at 4 months, followed by a decline between 7 and 11 months. This pattern is proposed to reflect infants’ initial sensitivity to slowly modulating coarse-grain prosodic information in the speech signal, which becomes progressively attuned to fine-grain information as infants acquire more extensive native language exposure. From this perspective, the strong delta-band tracking may support the perceptual organization of the continuous speech into larger prosodic units, providing a temporal scaffold for early speech processing. Consequently, as infants grow, speech processing may progressively rely more on faster neural oscillations, such as theta, facilitating the segmentation and encoding of meaningful linguistic units from continuous speech (Menn et al., 2023; Bosseler et al., 2013).

There is evidence that individual differences in the efficiency of cortical tracking of speech during the first months of life can be related to subsequent language development. Within this framework, phase synchronization, that is, the alignment of neural oscillatory phase with rhythmic input, at the stressed syllable rate of speech and drum beat rhythms at 2 months was predictive of children’s later vocabulary outcomes at 24 months (Ni Choisdealbha et al., 2023). Infants’ cortical tracking at the syllable but not stressed syllable rate at 11 months was also found to predict receptive and productive vocabulary at 24 months of age (Attaheri et al., 2024). In line with these findings, Menn et al. (2022) found that 10-months-old infants’ cortical tracking at the stressed syllable rate predicted receptive and productive vocabularies at 24 months. Studies in children also suggest that the tracking of speech rhythms is affected in children with and at risk of dyslexia. Di Liberto et al. (2018) found that children with dyslexia showed atypical cortical tracking when compared with age-matched and reading-level matched peers. Importantly, these results correlated with phonological awareness, a foundational skill for reading acquisition, and are consistent with studies in adults showing diminished sensitivity to speech edges in the amplitude envelope (Mandke, 2023) and reduced cortical tracking in dyslexic populations (Molinaro et al., 2016).

The association between early cortical tracking and subsequent language outcomes suggests that factors that facilitate cortical tracking may play a role in supporting language development. Consistent with this possibility, stimulus characteristics can modulate cortical tracking efficiency in young infants. Specifically, infants in the first year of life show stronger tracking of infant-directed speech compared to adult-directed speech (Kalashnikova et al., 2018, under review). Infant-directed speech is the speech typically used in adult-infant interactions, and it is characterized by higher and more variable pitch, slower tempo, exaggerated speech sounds, and more regular rhythmic properties compared to adult-directed speech (Trainor et al., 2000). Menn and colleagues (2022) examined the components in IDS that could facilitate infants’ cortical tracking by focusing on the acoustic adaptations at the syllable and stressed syllable rates. In their study, they tested German-learning 9-month-old infants’ cortical tracking to speech produced during natural mother-infant interactions. Their results showed that while infants efficiently tracked both IDS and ADS regardless of speech rate, cortical tracking at the stressed syllable rate was significantly stronger for IDS.

These findings suggest that rhythmic modulations present in natural IDS boost cortical tracking efficiency in young infants. Studies with adult participants provide more direct evidence for the effects of rhythmic modulation on cortical tracking efficiency, and that these effects emerge after brief rhythmic exposure and at the level of individual utterances. Falk et al. (2017) demonstrated that brief exposure to rhythmically-regular musical sequences led to significantly enhanced cortical tracking of subsequent speech. Specifically, they presented French native listeners with musical sequences that either reflected both the prosodic and syllabic structure of the French speech sequences or that followed an irregular rhythm. Participants showed greater cortical tracking of speech that followed the rhythmical musical sequences that reflected the structure of speech compared to the rhythmically irregular sequences. Fernández-Merino et al. (2025) expanded these findings by testing directly whether the positive modulation effects were due to the rhythmic regularity of the music sequences or their resemblance to the structure of speech used in the task. Spanish-Basque bilingual adults were presented with the same conditions as in Falk et al., with an additional third condition, where musical sequences maintained a regular rhythm reflecting the prosodic structure of speech but without matching its syllabic structure. Results showed successful modulation of cortical tracking in the presence of rhythmic regularity, regardless of whether or not the musical sequences matched the syllabic structure of speech. Interestingly, the bilingual participants were presented with the task in their two native languages, and language-specific effects emerged. Specifically, there was stronger delta-tracking for Spanish stimuli, but stronger delta- and theta-tracking for Basque stimuli, suggesting that the effects of rhythmic priming on cortical tracking may be shaped by language-specific rhythmic properties and listeners’ experience with the prosodic patterns of each language.

Previous work demonstrates that cortical tracking can be modulated through rhythmic priming in adults (Falk et al., 2017; Fernandez-Merino et al., 2025). However, it remains unclear whether similar modulation effects can be achieved in young infants, whose neural mechanisms for speech processing are still being attuned to the rhythmic properties of their native language, and who have more limited access to the linguistic content of the speech signal. While there is no direct evidence that brief priming enhances cortical tracking in infancy, previous work suggests that structured rhythmic input can shape language-related processing and communicative development. For example, Gerry, Unrau, and Trainor (2012) examined the effects of six months of musical training in infants who were 6 months old at the start of the study. Infants were randomly assigned to active musical classes, passive musical classes, or no musical training. Infants in the active musical training group demonstrated enhanced development of prelinguistic communicative gestures and social behaviors compared to the other groups, suggesting that actively engaging with musical rhythm in early infancy can support aspects of communicative development. Zhao and Kuhl (2016) also showed that 9-month-old infants benefited from musical exposure. In this study, infants from a control group and infants who had experienced one month of musical training were tested on their ability to detect temporal structure violations in both music and speech. With a shorter training experience than in Gerry et al. (2012), infants from the musical training group had an enhanced neural processing of temporal structure, shown by larger mismatch responses (MMR) to structure violations in speech and music. Similarly, beat and meter tracking is enhanced in infants enrolled in music classes and infants with musically trained parents (Cirelli et al., 2016, Flaten et al., 2022, 2025). In older children, rhythmic priming studies have further shown that brief exposure to regular musical rhythms can facilitate subsequent grammatical processing (e.g., Fiveash et al., 2020), supporting the idea that rhythmic structure may transiently enhance language processing mechanisms. Together, this evidence indicates that rhythmic experience throughout development can modulate neural sensitivity to temporal structure in both music and speech. Whether brief rhythmic priming that mirrors the temporal structure of speech can similarly enhance infants’ cortical tracking of speech remains unknown.

To address this question, we presented young infants with two speech conditions adapted from Fernández-Merino et al. (2025): a Regular condition (Matching-Regular in Fernández-Merino et al., 2025), in which speech was preceded by a regular musical sequence designed to mirror the temporal structure of the speech signal, and an Irregular condition, in which speech was preceded by a rhythmically irregular musical sequence. Importantly, infants completed this task at 6 and 10 months, so this longitudinal design allowed us to test potential developmental effects on the effectiveness of this rhythmic modulation. As it was the case for the adult participants in Fernández-Merino et al., (2025), the infants included in this study were acquiring Spanish and Basque in a bilingual context, allowing for an additional test of this rhythmic modulation across two typologically-different languages, and for its relation to individual differences in the amount of exposure to each language.

Based on the previous findings with adults, we anticipate a main effect of rhythmic condition in the delta and theta frequency bands: across both age groups, infants should show stronger cortical tracking when speech is preceded by Regular musical sequences compared to Irregular ones. This would confirm that exposure to regular, native-like speech rhythms generally facilitates cortical tracking of subsequent speech, as it has been observed for adults (Falk et al., 2017; Fernández-Merino et al., 2025). Second, we expect developmental changes in speech processing to manifest differently across frequency bands. In the delta band (∼2 Hz), which indexes prosodic processing, we predict robust tracking at both 6 and 10 months, with a greater effect of rhythmic priming (Regular vs Irregular condition) at 6 months, anticipating that infants would rely more on prosodic cues at this age, following previous literature (Attaheri et al., 2024). By 10 months, we predict a stronger priming effect in the theta band, associated with syllabic tracking, as infants begin to shift their attention toward finer-grained rhythmic cues.

## 2. Methods

### 2.1 Participants

A total of 46 Basque-Spanish bilingual infants (29 male, *M* age: 183 days, *SD* = 14.46) were recruited for the study and took part in the first session at 6 months. Infants were recruited through the Basque Center on Cognition, Brain, and Language (BCBL) Babylab database. Nine infants were excluded from the analysis due to (1) fussiness (N = 3) and (2) failure to comply with the language exposure criteria for bilingualism (N = 6), so they were not invited for the second session. When infants turned 10 months, they were invited to come back to the lab for a second session, and 32 infants (17 male, *M* age: 317 days, *SD* = 13.95) did so. One infant was excluded due to a technical error. Thus, the final analyses included data from 37 infants for the first session at 6 months, and 31 infants for the second session at 10 months.

All infants were born after at least 37 weeks of gestation and with no apparent health problems. The Ethics and Scientific Committee of the BCBL, following the declaration of Helsinki, approved the study protocol (approval number 150621SM). Parents were informed about the study and provided informed consent, and infants were given a diploma and a souvenir for their participation.

### 2.2 Participants language background

Infants’ language background was estimated based on a parental Language Exposure Questionnaire (Bosch & Sebastián-Gallés, 1997, Molnar et al., 2014). This questionnaire gives an estimate of infants’ language exposure patterns to each of the languages from birth until the date of testing. Infants were considered to be bilingual if they received a maximum of 75% and a minimum of 25% exposure to each of their languages and no more than 10% of exposure to a third language (Pearson, Fernández, Lewedeg, & Oller, 1997). At 6 months, infants were exposed to Spanish on average 54.76 % of the time (SD = 14.29, range = 27.3–75.1) and to Basque 45.23 % of the time (SD = 14.29, range = 24.9–72.7). At 10 months, infants received 53.66% exposure to Spanish (SD = 14.33, range = 27.4–74.9) and 46.33% exposure to Basque (SD = 14.33, range = 25.1–72.6). All infants received exposure to their two languages from their parents at home and from regular interactions with close relatives or other members of the community. Six of 37 infants at 6 months and 5 of 31 infants at 10 months heard one language from each parent. All other infants heard both languages from both parents.

Previous studies measuring language dominance in bilingual infants have established infants’ dominant (L1) and nondominant (L2) language based on the proportion of exposure to each language (e.g., Kalashnikova et al., 2022; Singh et al., 2023), whereby the language with >50% of exposure is typically considered dominant. Following this criterion, there were a total of 27 Spanish and 11 Basque dominant infants at 6 months, and a total of 21 Spanish and 11 Basque dominant infants at 10 months. Importantly, the language exposure questionnaire captures cumulative exposure patterns from birth to the time of testing, including both infant-directed and overheard speech. Thus, it is noteworthy that while the estimates obtained from these questionnaires are highly effective at quantifying infants’ relative accumulated exposure to each language during their lifetime, they are limited at reflecting more recent changes in infants’ language dominance patterns. For instance, an infant who received more exposure to Spanish than Basque until 6 months and who then entered Basque-medium day-care would have received more exposure to Spanish than Basque overall (a common situation in our community), but their dominance may be in the process of shifting to Basque at the time of testing. Thus, to better capture infants’ emerging language knowledge, we complemented exposure-based estimates with a measure of functional dominance based on vocabulary size, defining dominance as the language in which infants were reported to understand and produce more words (Byers-Heinlein et al., 2024). For this purpose, we administered the Spanish and Basque versions of the MacArthur–Bates Communicative Development Inventories (Words and Sentences CDI; Barreña et al., 2008; Jackson-Maldonado et al., 2013) when infants were 10 and 18 months of age. These data were available for 24 out of the 31 infants with 10-month data (the remaining families did not return the CDI). Notably, 21 of these 24 infants were reported to understand and produce more words in Basque than in Spanish.

### 2.3 Musical experience

Caregivers also completed a short questionnaire about infants’ musical background, based on the Music@Home questionnaire (Politimou et al., 2018) and adapted to the specifics of our study. Overall, infants had minimal formal musical exposure, and musical experience did not differ across age or language groups. Further information about this measure is provided in the Supplementary Materials.

### 2.4 Stimuli

Stimuli were the same used in Fernández-Merino et al., (2025) and are available at https://osf.io/r4nz9/?view_only=8dab2eb264214abd8d31d555c2e338b7. Speech stimuli consisted of 64 12-syllable sentences in Spanish and 64 13-syllable sentences in Basque. In order to ensure natural and highly rhythmic stimuli that highlighted the syllable information and because the same stimuli were used in an identical study with infants, all stimuli were recorded by a female Basque-Spanish bilingual speaker in IDS, since it provides the ideal context to stress and highlight the temporal modulations of speech (Leong et al., 2014).

This experimental design required two types of musical stimuli to be presented in the Regular and Irregular conditions. The musical stimuli for the Regular condition were created based on the temporal structure of the sentences, as it reflected the temporal structure and pitch contour of linguistic stimuli. For this purpose, we extracted the rhythmic structure of the sentences to create the beat structure. This structure was introduced in Musescore 3 (https://musescore.org/es), an open-source software where, using a grand piano timber, we created the melodic contour from the pitch contour of the sentences. The Irregular musical sequences were created by altering the pitch and note duration of the Regular sequences, thus also altering their regular meter. The melodic sequences that comprised a condition formed a musical structure of A A A A’ (A’: slight variation of the melodic contour and temporal structure of A), following the common musical pattern of Basque and Spanish infant-directed songs.

The speech sequences and the melodic sequences had a duration of 2500 msec in the Spanish blocks, and 3750 msec in the Basque blocks. To ensure highly rhythmic stimuli, each speech sequence was cued by a metronome at regular rate of 120 bpms in Spanish and 100 bpm in Basque, reflecting the rate of stressed syllables, obtaining a regular meter with an inter-onset interval between accented syllables of 500 msec in Spanish and 625 in Basque. The musical sequences were created based on the temporal patterns of the sentences. There was a constant inter-onset interval between accented beats for the Regular musical sequences, and no constant inter-onset interval for the Irregular sequences.

Consistent with the notion that Spanish and Basque have different rhythmic properties (Molnar et al., 2016, Molnar et al., 2014), spectral analyses of the melodic and speech sequences revealed different frequencies for each language, with averaged peaks at 2 Hz and 4 Hz in Spanish and peaks at 1.6 Hz and 4 Hz in Basque. The corresponding figure showing these analyses can be found in the Supplementary Materials. Besides, we estimated the syllable rate of the sentences using an automatic algorithm based on the method from De Jong & Wempe (2009) and Praat (Boersma & Weenink, 2021). The overall stressed rate in Spanish was found to be 2.15 Hz (SD = 0.25) and 1.56 Hz (SD = 0.28) in Basque, with the average syllable rate in Spanish being 4.23 Hz (SD = 0.33) and 4.16 Hz (SD = 0.28) in Basque, showing a high degree of similarity between the two languages. Based on the frequency resolution of 0.4 Hz used in our speech-brain coherence analysis, the frequency bin that most closely aligned with the stressed syllable rate was 2 Hz for Spanish and 1.6 Hz for Basque. The syllabic rate in both languages was most strongly represented at 4 Hz. These peaks were expected because they correspond to the prosodic (reflected in the delta frequency band, ∼2Hz) and syllabic (reflected in the theta frequency band, ∼4Hz) rhythms. No specific frequency was reflected in the Irregular musical sequences.

### 2.5 Procedure

A graphical example of the procedure is illustrated in Figure 1. Participants completed two conditions in Spanish and Basque in a single experimental session at 6 and 10 months of age: speech preceded by Regular musical cues and speech preceded by Irregular musical cues. Stimulus presentation was structured in 16 blocks of four trials. Only one type of condition, Regular or Irregular, and one language, Spanish or Basque, was used within a single experimental block. Blocks were grouped by language such that infants were presented with all blocks from one language before switching to the other language (e.g., the full Spanish portion followed by the full Basque portion, or vice versa), with language order counterbalanced across participants. There was a total of 64 blocks, with a total experiment duration of ∼ 22 minutes. There were 16 blocks per language (Basque or Spanish) and per rhythm condition (Regular and Irregular), each condition had a duration of ∼ 5’ 30". All linguistic stimuli were preceded by Regular and Irregular musical sequences. A single block consisted of four trials and had a total duration of 20 sec in the Spanish blocks and 30 sec in the Basque blocks. Each trial had a duration of 5000 msec in the case of Spanish blocks and 7500 msec in the case of Basque blocks and comprised one melodic sequence immediately followed by a simple sentence. The stimulus onset asynchrony (SOA) between the melodic sequence and the sentence was always 500 msec for the Spanish trials and 625 msec for the Basque trials.

**Figure 1.**
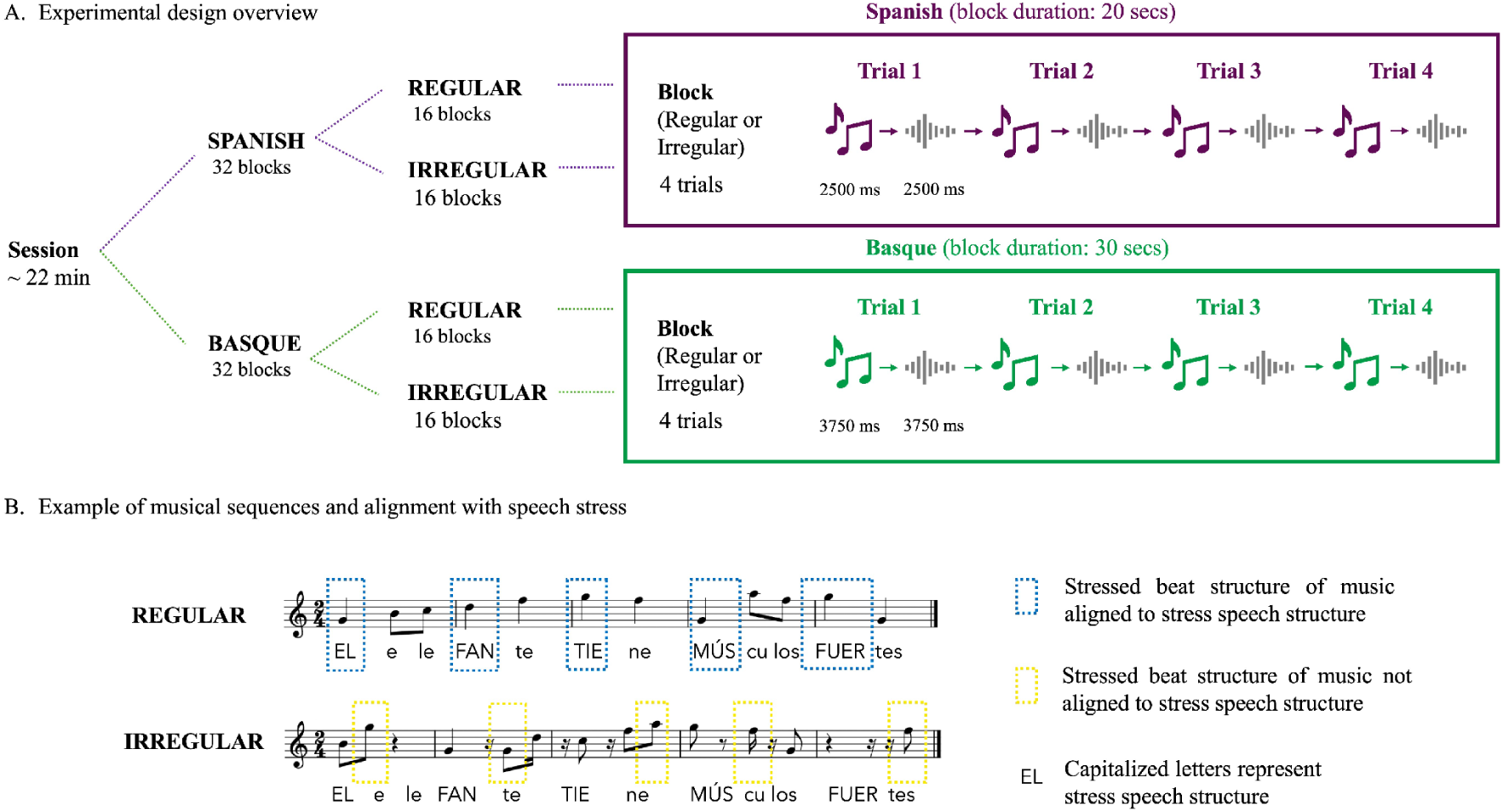
Experimental design and stimuli. **(A)** Overview of the experimental design. Infants completed the task at 6 and 10 months of age, and were presented with the Spanish and Basque blocks. For each language, infants were presented with 32 blocks, comprising 16 Regular and 16 Irregular blocks. Each block had four trials and used only one rhythmic condition throughout. In each trial, a musical sequence was followed by a sentence. **(B)** Examples of the musical sequences used in the Regular and Irregular conditions and their relation to the stress structure of the subsequent speech. In the Regular condition, stressed beats in the musical sequence were aligned with stressed syllables in the sentence. In the Irregular condition, stressed beats did not align with stressed syllables. Capitalized letters indicate stressed syllables in the speech sequence (example sentence: El elefante tiene músculos fuertes; English translation: The elephant has strong muscles). Blue dashed boxes indicate alignment between musical beats and stressed syllables, whereas yellow dashed boxes indicate a lack of alignment.

The order of condition and language was counterbalanced across participants. Stimuli were presented in PsychoPy (v.1.80.04) via two speakers positioned about one meter in front of the participants, at 65 dB. Participants were in front of a screen showing cartoons to entertain them, while a research assistant entertained them with toys as well.

### 2.6 EEG data acquisition and preprocessing

EEG data were acquired using a BrainAmp amplifier and BrainVision Recorder software (Brain Products, Germany). EEG was recorded using 16 active electrodes that were positioned according to the international 10–20 system (Jasper, 1958): F3, Fz, F4, FC1, FC2, T7, C3, Cz, C4, T8, CP1, CP2, P3, Pz, P4, M2. Scalp-electrode impedance was kept below 20 kΩ for scalp electrodes to ensure high-quality EEG recordings. EEG data were sampled at a rate of 1000 Hz and band-pass filtered online from 0.1 to 1000 Hz. The left mastoid served as the online reference, and electrode FCz was used as the ground.

All the following processing and analysis steps were implemented in Matlab (The MathWorks, Inc., Natick, US) using FieldTrip (Oostenveld et al., 2011) and available custom code (Fernández-Merino & Lizarazu, 2023). Movement artifacts were detected and removed using iMARA (Haresign et al., 2021). Data from the Spanish blocks were segmented from 0 s to 2.5 sec relative to stimulus onset. Data from the Basque blocks were segmented from 0 s to 3.75 sec after the onset of each trial. Since the trial length in the Basque blocks (3750 msec) led to a frequency resolution of ∼0.26 Hz, we created new trials of 2500 msec with an overlapping window of 1250 msec to achieve a frequency resolution of ∼0.4 Hz for the analyses, to match the Spanish blocks. These trials were re-referenced off-line to the average activity of the two mastoids and low pass filtered below 30 Hz, since we did not expect any coherence effect above this threshold (see Gross et al., 2013). The remaining trials were visually inspected. A minimum of 60% artifact-free trials per participant was required for inclusion in subsequent analyses.

## 3. Analysis

### 3.1 Speech and music brain coherence

We quantified the phase synchronization (referred to as coherence hereafter) between the brain signal and the corresponding auditory stimuli envelopes weighted by their relative amplitude using Hilbert coherence over time (Fernández-Merino & Lizarazu, 2023; Molinaro et al., 2016). Broadband amplitude envelope of the audio signals was obtained from the Hilbert transformed broadband stimulus waveform (Drullman et al., 1994). Then, audio envelopes were resampled from 44100 Hz to 1000 Hz. For each experimental condition, coherence between the artifact free epochs and the corresponding audio envelope was calculated in the 0.4 – 10 Hz frequency band with 0.4 Hz frequency resolution (inverse of the trial duration). Following this procedure, we obtained a coherence value for each (i) participant, (ii) condition, (iii) EEG sensor and (iv) frequency bin below 10 Hz.

For the subsequent analyses, we extracted the maximum coherence values across both frequency bins and channels, separately for each participant and condition. In the Spanish blocks, maximum values were extracted from 1.6 Hz to 2.4 Hz in the case of delta, and 3.6 Hz to 4.4 Hz in the case of theta. In the Basque blocks, maximum values were extracted from 1.2 Hz to 2 Hz in the case of delta, and 3.6 Hz to 4.4 Hz in the case of theta.

## 4. Results

### 4.1 Statistical analysis

We conducted separate analyses for each language version of the experiment (Spanish and Basque), stimulus type (musical sequences and speech sequences), and frequency band of interest (delta and theta). Linear mixed-effects models were fitted with coherence values as the dependent variable and a two-way interaction between Condition (Regular, Irregular) and Age (6, 10 months) as fixed effects. Random intercepts were specified for participants. Continuous variables were scaled and centered around zero to facilitate model interpretation. Main effects and interactions were further explored using Bonferroni-corrected post hoc comparisons. Analyses were conducted in R using the lme4 (Bates, 2018) and lmerTest (Kuznetsova et al., 2017) packages.

Prior to assessing whether exposure to musical sequences with different rhythmic properties modulated cortical tracking of subsequent speech, we first examined cortical tracking of the musical sequences themselves. Although these analyses were not central to our primary hypotheses, we expected Regular musical sequences, because of their greater temporal predictability and rhythmic structure, to elicit stronger cortical tracking than Irregular sequences. This served as a manipulation check to confirm that the rhythmic regularity of the musical input was effective before evaluating its influence on speech processing. Accordingly, the results are organized into analyses of cortical tracking to musical sequences and cortical tracking to speech sequences, each presented separately for Spanish and Basque. Results for musical sequences are shown in Figure 2 and results for speech sequences are shown in Figure 3. Correlational analyses examining associations between cortical tracking and musical experience questionnaire measures yielded no significant effects and are therefore reported in the Supplementary Materials.

**Figure 2:**
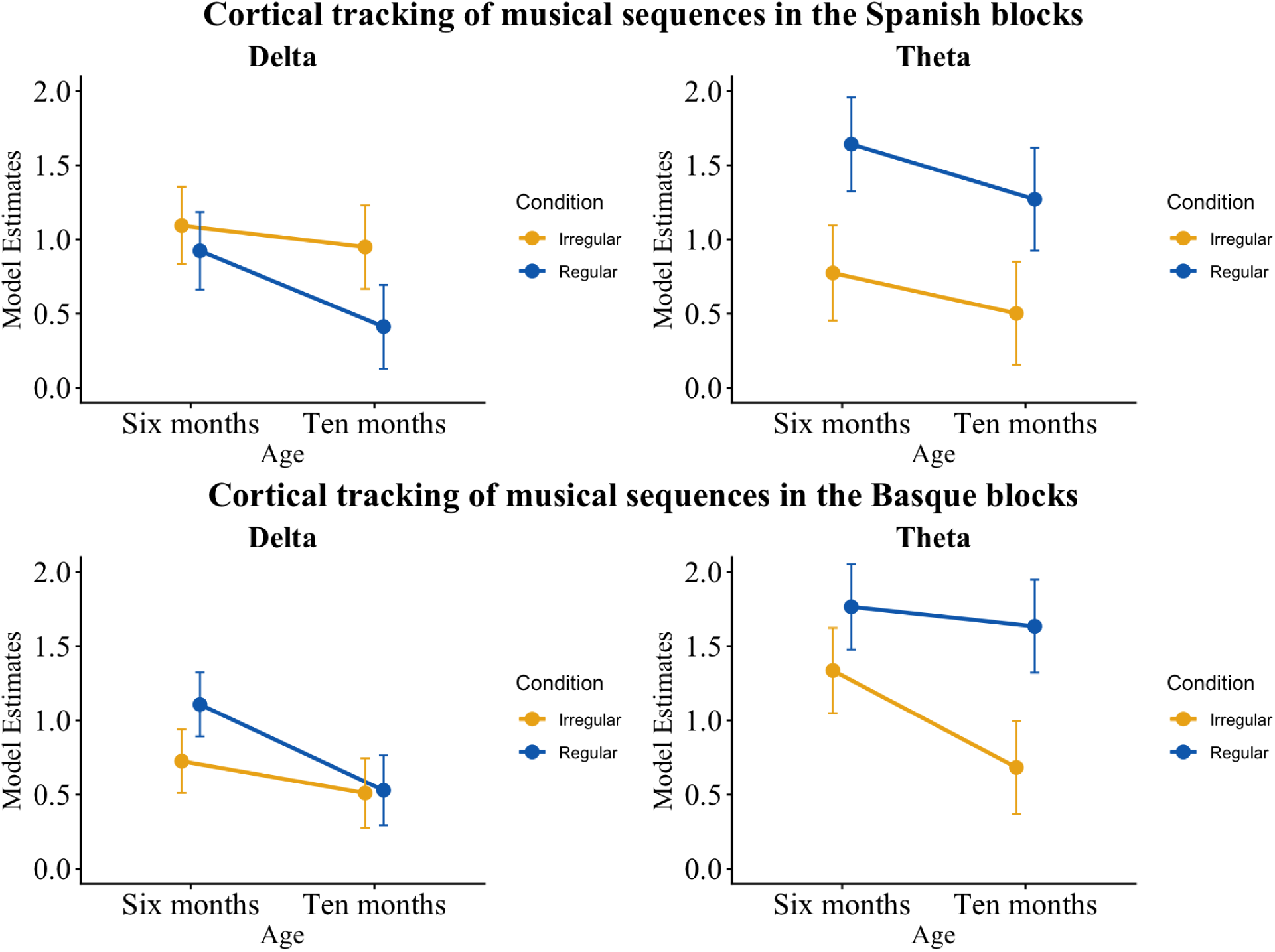
Overview of model estimates of cortical tracking to musical sequences. Top panel shows cortical tracking to musical sequences in the Spanish blocks. Bottom panel shows cortical tracking to musical sequences in the Basque blocks. Error bars show standard error from the mean.

**Figure 3.**
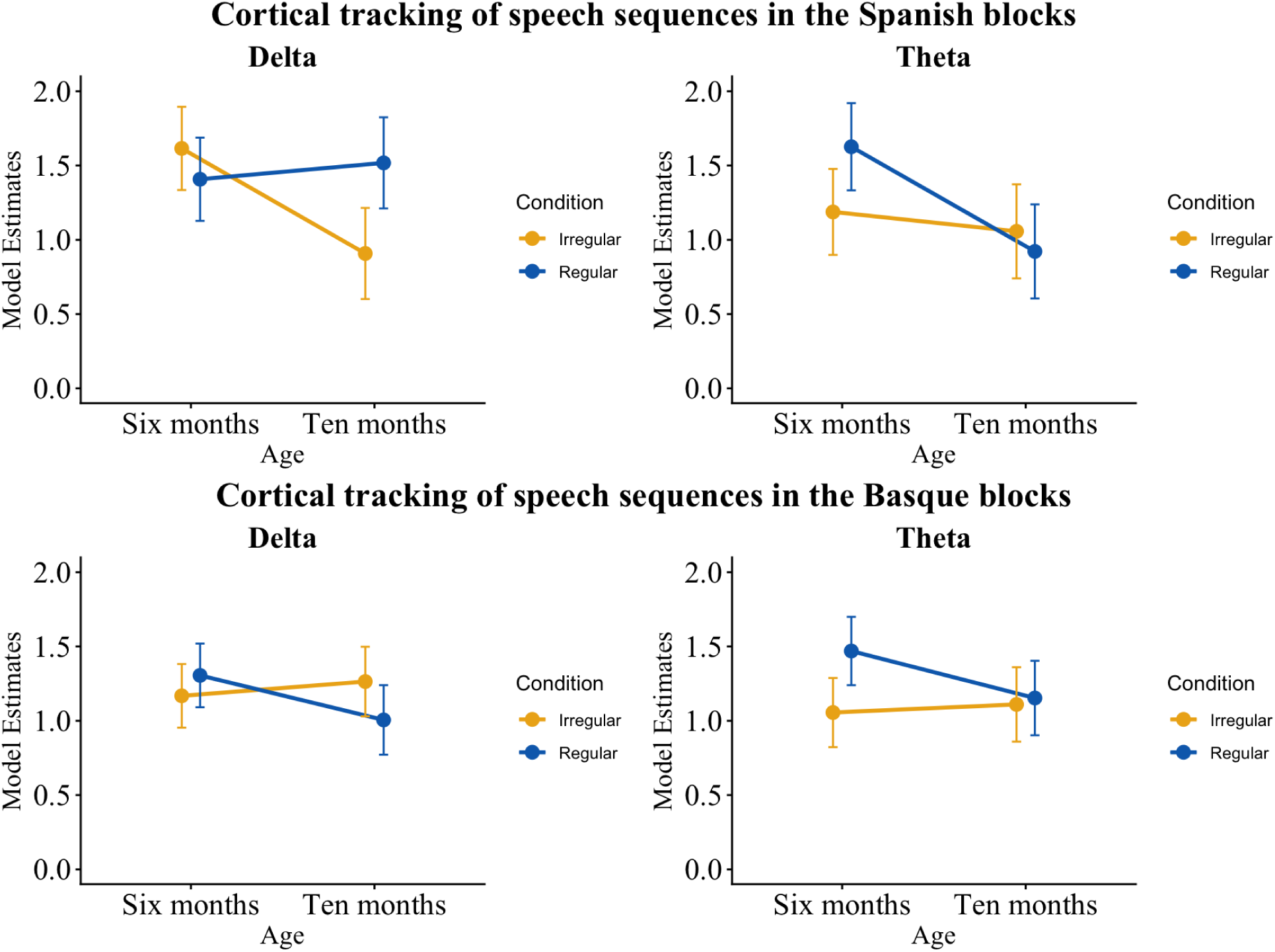
Overview of model estimates of cortical tracking of speech sequences. Top panel shows cortical tracking to speech sequences in the Spanish blocks. Bottom panel shows cortical tracking to speech sequences in the Basque blocks. Error bars show standard error from the mean.

### 4.2 Musical sequences

#### 4.2.1 Cortical tracking to musical sequences in Spanish blocks

For the delta band, the model revealed a main effect of Condition (*F* (1, 94) = 7.446, *p* <.01), a main effect of Age (*F* (1, 104) = 6.287, *p* = .013), and no significant Condition by Age interaction (*F* (1, 94) = 1.988, *p* = .16). Post Hoc comparisons revealed that coherence was significantly higher in the Irregular condition compared to the Regular condition (*β* = 0.353, *SE* = 0.13, *t* = 2.729, *pbonf* < .001), and higher at 6 months compared to 10 months (*β* = 0.328, *SE* = 0.131, *t* = 2.501, *pbonf* = .013).

In the theta band, the model also revealed a main effect of Condition (*F* (1, 97) = 26.079, *p* < .001), and Age (*F* (1, 106) = 3.974, *p* = .04), with no significant Condition by Age interaction (*F* (1, 97) = 0.095, *p* = .758). However, contrary to the delta band, coherence was significantly higher in the Regular condition compared to the Irregular condition (*β* = −0.818, *SE* = 0.16, *t* = −5.106, *pbonf* < .0001). Coherence was again higher at 6 months compared to 10 months (*β* = 0.322, *SE* = 0.162, *t* = 1.989, *pbonf* = .04).

#### 4.2.2 Cortical tracking to musical sequences in the Basque blocks

In the delta band, the model revealed no main effect of Condition (*F* (1, 96) = 3.151, *p* = .079), a main effect of Age (*F* (1, 105) = 12.366, *p* <.001), and no significant interaction (*F* (1, 96) = 2.588, *p* = .11). Like the Spanish blocks, coherence was significantly higher at 6 months compared to 10 months (*β* = 0.397, *SE* = 0.113, *t* = 3.51, *pbonf* < .001).

In the theta band, the model revealed a main effect of Condition (*F* (1, 96) = 26.311, *p* < .001), a main effect of Age (*F* (1, 103) = 8.228, *p* < .01), and a marginal Condition by Age interaction (*F* (1, 96) = 3.755, *p* = .055). Post Hoc comparisons revealed that coherence in the Regular condition was significantly higher than in the Irregular condition (*β* = −0.69, *SE* = 0.134, *t* = −5.129, *pbonf* <.001) and significantly higher at 6 months compared to 10 months (*β* = 0.392, *SE* = 0.137, *t* = 2.862, *pbonf* < .01).

### 4.3 Speech sequences

#### 4.3.1 Cortical tracking to speech sequences in the Spanish blocks

In the delta band, the model revealed no main effect of Condition (*F* (1, 95) = 1.937, *p* = .167), a main effect of Age (*F* (1, 104) = 4.22, *p* = .04), and a significant Condition by Age interaction (*F* (1, 95) = 7.991, *p* < .01). Coherence was significantly higher at 6 months compared to 10 months (PostHoc: *β* = 0.299, *SE* = 0.146, *t* = 2.052, *pbonf* = .04). Coherence in the Regular condition was significantly higher than in the Irregular condition at 10 months (*β* = −0.61, *SE* = 0.213, *t* = −2.86, *pbonf* = .03) but not at 6 months (*β* = 0.207, *SE* = 0.195, *t* = 1.063, *pbonf* = 1).

In the theta band, there was no main effect of Condition (*F* (1, 96) = 1.03, *p* = .319), but a main effect of Age (*F* (1, 105) = 7.548, *p* < .01), and no significant Condition by Age interaction (*F* (1, 96) = 3.578, *p* = .06). Coherence was again significantly higher at 6 months compared to 10 months (*β* = 0.418, *SE* = 0.152, *t* = 2.742, *pbonf* < .01).

#### 4.3.2 Cortical tracking to speech sequences in the Basque blocks

In the delta band, we found no main effects of Condition (*F* (1, 96) = 0.312, *p* = .577), Age (*F* (1, 105) = 0.873, *p* = .352), nor significant interaction (*F* (1, 97) = 3.34, *p* = .07).

In the theta band, we found a main effect of Condition (*F* (1, 95) = 4.161, *p* = .04), with coherence in the Regular condition being significantly higher than in the Irregular condition (*β* = −0.229, *SE* = 0.112, *t* = −2.04, *pbonf* = .044). There was no main effect of Age (*F* (1, 103) = 1.334, *p* = .25), or significant interaction (*F* (1, 95) = 2.737, *p* = .101).

## 5. Discussion

This study investigated whether immediate exposure to musical rhythms that resemble the temporal structure of speech can modulate infants’ cortical tracking of speech, and how such effects evolve between 6 and 10 months of age. To do so, we presented infants with sentences in their two native languages, Spanish and Basque, that were preceded either by regular musical sequences (matching prosodic and syllabic structure of the speech sequences) or by irregular sequences. We hypothesized that (1) infants would show enhanced cortical tracking in the Regular condition compared to the Irregular condition, reflecting how exposure to temporally structured musical input facilitates cortical tracking of speech; and (2) these effects would vary as a function of age, reflecting developmental changes in delta versus theta cortical tracking mechanisms.

Across languages, our findings suggest that even brief rhythmic experience can dynamically shape infants’ neural encoding of speech. Specifically, exposure to Regular musical sequences modulated infants’ cortical tracking of subsequent speech, mirroring previous findings with adults that even brief rhythmic input can influence how speech is processed (Falk et al., 2017; Fernández-Merino et al., 2025). Importantly, the effect of rhythmic priming was not uniform across frequency bands or languages. Instead, we show that rhythmic musical input selectively modulates speech tracking at linguistically relevant temporal scales, suggesting that neural oscillations exploit rhythmic regularities in the environment to optimize speech processing. In the Spanish blocks, when speech was preceded by musical sequences that mirrored native rhythmic patterns (the Regular condition), cortical tracking in the delta band (∼2 Hz) was significantly enhanced, but only at 10 months, not at 6 months. In contrast, for Basque, the enhancement occurred in the theta band (∼4 Hz) rather than in the delta frequency band..

There are several factors that could account for these findings. First, this effect could reflect inherent differences in language rhythmic structure: Spanish is predominantly syllable-timed with relatively stable prosodic patterns, while Basque exhibits mixed rhythmic properties (Aurrekoetxea et al., 2013; Molnar et al., 2016). In this view, regular rhythms could selectively enhance speech tracking at the temporal scale most relevant for each language, suggesting that rhythmic priming operates by aligning neural oscillations to linguistically meaningful temporal structure rather than by producing a general increase in cortical tracking, irrespective of the language. More broadly, these findings align with theoretical frameworks proposing that neural oscillations act as active temporal prediction mechanisms during speech processing (Morillon et al., 2015; Giraud & Poeppel, 2012). Our findings suggest that such predictive alignment mechanisms are already operational in infancy and remain flexible to short-term rhythmic experience. Second, the differences between stressed syllable (∼2 Hz, delta) and syllable (∼4 Hz, theta) rate effects could emerge from the effects of infants’ language development and maturation of oscillatory neural activity in the delta and theta frequency bands. As stated in the Introduction, delta tracking is associated with prosodic grouping and slower speech temporal structure, whereas theta tracking supports syllabic parsing and finer-grained segmentation. We argue that infants show a developmental shift from broad sensitivity to slow rhythmic structure toward more selective tuning to faster, linguistically relevant timescales, as shown in recent infant cortical tracking literature (Attaheri et al., 2024; Menn et al., 2023). The developmental differences observed here may also reflect broader maturational changes in large-scale cortical networks supporting temporal integration (Háden et al., 2025; Ortiz-Barajas et al., 2023). Early in development, infants may rely more heavily on slower oscillatory sampling strategies that support broad prosodic processing, whereas increasing specialization of theta band activity could reflect emerging efficiency in segmentation and syllabic encoding. This interpretation is consistent with proposals that maturation of oscillatory hierarchies constitutes a foundational mechanism for the development of speech tracking and language development (Menn et al. 2023; Giraud et al., 2012).

While our results are consistent with this framework, they cannot be explained without considering the role of infants’ language-specific exposure in our study. In our sample, infants’ degree of exposure to each language was likely changing over time. Although cumulative exposure questionnaires classified many infants as Spanish-dominant, vocabulary size data indicated that many infants developed stronger Basque vocabularies. This discrepancy reflects the dynamic nature of bilingual input: cumulative exposure from birth may not capture recent shifts in infants’ functional language experience (e.g., entry into Basque-only daycare). Thus, we propose that the language-specific effects observed here are likely to arise from rhythmic priming interacting with infants’ current language-specific competence and with each language’s temporal structure. This interpretation is consistent with previous findings showing that cortical tracking in Spanish-Basque bilinguals is modulated by language exposure (Pérez-Navarro et al., 2024). Importantly, our sample also provides information about studying rhythmic adaptation in bilingual infants, who must flexibly track multiple prosodic systems simultaneously. From this perspective, bilingual experience may place increased demands on the mechanism of cortical tracking, possibly amplifying sensitivity to rhythmic cues in the environment. Regular musical sequences would then act as a temporal scaffold, aligning neural oscillations to the prosodic or syllabic rhythms that are most informative for processing the upcoming speech signal in that language. However, this interpretation should be directly tested in future work comparing bilingual and monolingual infants, as well as examining how variability in language exposure shapes neural tracking of speech across development. While we cannot exclude the possibility that the bilingual status of the participants contributed to the observed patterns, the present study was not designed to test this question directly.

Taken together, our results indicate that brief exposure to temporally structured input can shape infants’ cortical tracking to subsequent speech, and that this effect depends on both the developmental stage and the rhythmic properties of the language being processed. Our findings support the view that music and speech share partially overlapping temporal processing mechanisms (Peretz et al., 2015; te Rietmolen et al., 2024). The observation that musical rhythms selectively modulate subsequent speech tracking suggests that the transfer effects observed here and in Fernández-Merino et al. (2025) arise from shared rather than domain-specific mechanisms, supporting a perspective where rhythm serves as a domain-general scaffold for auditory processing (Fiveash et al., 2021). In that regard, these findings raise an important mechanistic question: what exactly is being primed by the musical sequences? If musical rhythms influence speech tracking by synchronizing neural oscillations to specific temporal patterns, then this synchronization should also be visible in how infants track the musical sequences themselves. Consistent with this idea, infants in this study generally showed stronger cortical tracking for regular than for irregular musical sequences, particularly in the theta band (∼4 Hz). We also observed an unexpected effect of condition in the delta band (∼2 Hz) in Spanish blocks, as coherence was higher for the Irregular than the Regular sequences. Although this finding deviates from the general pattern observed in our study, it is not unprecedented: similar responses to irregular input have been reported in adult participants (Falk et al., 2017; Fiveash et al., 2020). One possible explanation is that irregular sequences, precisely because they lack overt rhythmic structure, require greater cognitive effort to process, leading to stronger low frequency tracking. Rather than passively synchronizing to a predictable beat, the brain may engage in active temporal prediction to detect patterns in irregular input, thereby enhancing tracking in the delta band. This sensitivity to irregular input may reflect early developmental mechanisms for detecting structure in complex auditory environments, where infants must learn to extract regularities even when they are not overtly marked. In all other musical conditions, including the theta band (∼4 Hz) for both Spanish and Basque blocks, coherence was higher for the Regular condition, consistent with prior work showing infants’ sensitivity to beat-based structure in music (Lenc et al., 2023; Flaten et al., 2022; Cirelli et al., 2016). It is important to note that more isochronous stimuli inherently elicit stronger coherence responses, particularly at beat-related frequencies, suggesting that some of the observed effects may be driven by the physical properties of the stimuli themselves. However, the modulation from musical rhythm to subsequent speech tracking suggests that the observed effects cannot be fully reduced to low-level acoustic regularity alone. Thus, it appears to be a specific and isolated deviation from the broader pattern, and it does not undermine our central interpretation of the speech findings.

### 5.1 Limitations and future directions

The present findings provide evidence that brief exposure to regular musical rhythms can modulate infants’ cortical tracking of speech. However, the design does not allow us to determine the extent to which this modulation reflects an enhancement relative to baseline speech processing, as we did not include a speech-only condition. Our goal was to compare the effects of rhythmically aligned and misaligned musical cues, building on previous studies of cortical tracking with and without musical exposure (e.g., Falk et al., 2017; Fernández-Merino et al., 2025; Molinaro et al., 2016; Molinaro & Lizarazu, 2018). Nevertheless, including a speech-only condition in future work would help establish whether regular rhythmic cues enhance cortical tracking above baseline levels or whether irregular cues disrupt neural synchronization to speech.

More broadly, the mechanisms underlying the observed modulation remain to be fully understood. The present study focused on rhythmic regularity, but the stimuli also contained other prosodic cues that are known to support speech processing in infancy. In particular, the speech materials were recorded in infant-directed speech, which is characterized by exaggerated prosodic structure and enhanced pitch contours (Fernald, 1984; Kuhl, 1988; Trainor, 2000). These characteristics were intentionally preserved because they reflect the speech infants encounter in natural interactions. However, this design does not allow us to determine the relative contributions of rhythmic and pitch-based cues to the observed effects. Future studies that independently manipulate rhythmic regularity and pitch contours could clarify how different prosodic dimensions interact to support cortical tracking during early development.

Finally, an important question is whether the short-term modulation of cortical tracking observed here has consequences for longer-term language development. Given the growing evidence linking cortical tracking to subsequent language outcomes, it will be important to determine whether repeated exposure to rhythmically structured input can produce lasting changes in speech processing or language learning. Addressing this question could help clarify the developmental significance of rhythmic modulation and its potential relevance for interventions aimed at supporting infants at risk for language difficulties.

## 6. Conclusion

This study provides new insights into how rhythmic exposure influences cortical tracking in infants. Our findings highlight the complex interplay between musical and linguistic rhythms, showing that immediate exposure to rhythmic structures resembling speech rhythms can modulate infants’ speech tracking, particularly at the stressed syllable rate (∼2 Hz) in Spanish and the syllable rate (∼4 Hz) in Basque. Besides, our findings suggest that cortical tracking in infancy is a dynamic mechanism that can be shaped by rhythmic context while it develops with age and language exposure. More broadly, these findings position rhythmic structure as a fundamental organizing principle of early language processing. By demonstrating that brief rhythmic experience dynamically shapes cortical tracking in infancy, we propose here that oscillatory tracking acts as a flexible and dynamic mechanism between temporal structure in the environment and language development.

## Data and code availability

All code used for data processing and analysis available at https://doi.org/10.6084/m9.figshare.22740353.v3. Data available upon request to the authors. Stimuli are available at https://osf.io/epa4n/?view_only=76177c6fe76b45fbb8eeb33e8446b048.

## Acknowledgements

We want to thank the participants for their volunteer contribution to our study. This work was supported by the Basque Government through the BERC 2022-2025 program and Funded by the Spanish State Research Agency through BCBL Severo Ochoa excellence accreditation CEX2020-001010/AEI/10.13039/501100011033. LFM’s work was supported by a Predoctoral Grant from the Spanish Ministry of Science, Innovation and Universities and the European Social Fund, PRE2019-087623. ML’s work was supported by the Ramón y Cajal Postdoctoral Fellowship from the Spanish Ministry of Science, Innovation and University (grant RYC2022-035497-I) and by the European Commission (European Research Council) (grant ERC-2023-ADG–101140427). NM and MK received support from the Spanish Ministry of Science, Innovation and University, (NM: grants RTI2018-096311-B-I00, PID2022-136991NB-I00, PCI2022-135031-2, AIA2025-163317-C33, PDC2025-166757-I00; MK: grants RYC2018-024284-I, PID2022-136986NB-I00).

## CRediT statement

Conceptualization: LFM, NM, MK; Data curation and Formal Analysis: LFM; Funding Acquisition: MK; Investigation and Methodology: LFM; Project Administration: LFM, MK; Software: LFM, ML; Supervision: NM, MK; Writing - original draft: LFM; Writing - review and editing: LFM, NM, MK.

