## Supplementary materials for "Modulating Neural Tracking of Speech in Infants"

#### 1. *Musical exposure measures*

During the first visit to the lab at 6 months, infants' musical exposure was measured using a musical background questionnaire created at our lab (see Appendix). This questionnaire was created by including and adapting questions from validated questionnaires previously used in infants and adults (Music@home, add reference, Gold-MSI, Parent Music Activities). The questionnaire was constituted by 15 items and collected information about parents' musical education, music listening in the home (frequency and genres), activities, and singing interactions with the infants (frequency; context, e.g., play, sleep time; type of songs, e.g., lullabies, play songs). Some of the questions were open and descriptive (e.g., concerning the types of songs used with the infants, or the types of instruments/objects/toys used to make musical sound). Three variables were extracted from this questionnaire: parents' musical background and beliefs, infants' rhythmical exposure and infants' reactions towards music evaluated on a 7-Likert scale and transformed to percentages for analysis. Only infants' rhythmical exposure was included as a variable for later analysis. Data was collected from all participants. Infants in this study were all regularly exposed to music at home, with at least 1 hour of infant-directed singing exposure from their mothers. From the full sample, 19 out of 36 infants were actively exposed to their parents playing instruments at home. No infant was attending musical classes at the time of collecting musical background data.

##### *1.1 Statistical analysis*

To analyze the relationship between cortical tracking and infants' musical exposure, we conducted correlation analyses between infants' cortical tracking values and infants' music exposure data. These consisted of correlation analysis between coherence values for each language in the stressed (~2 Hz) and in the syllable rate (~4 Hz) in the Regular condition, and infants' individual musical experience scores.

#### *1.2 Relationship between infants' cortical tracking and infants' musical exposure*

We ran correlations between infants' cortical tracking to the speech and musical sequences in the Regular condition in the stressed syllable and syllable rate and infants' rhythmical exposure, shown in Figure 1. We found no significant correlations between infants' cortical tracking of speech and musical sequences and their music exposure.

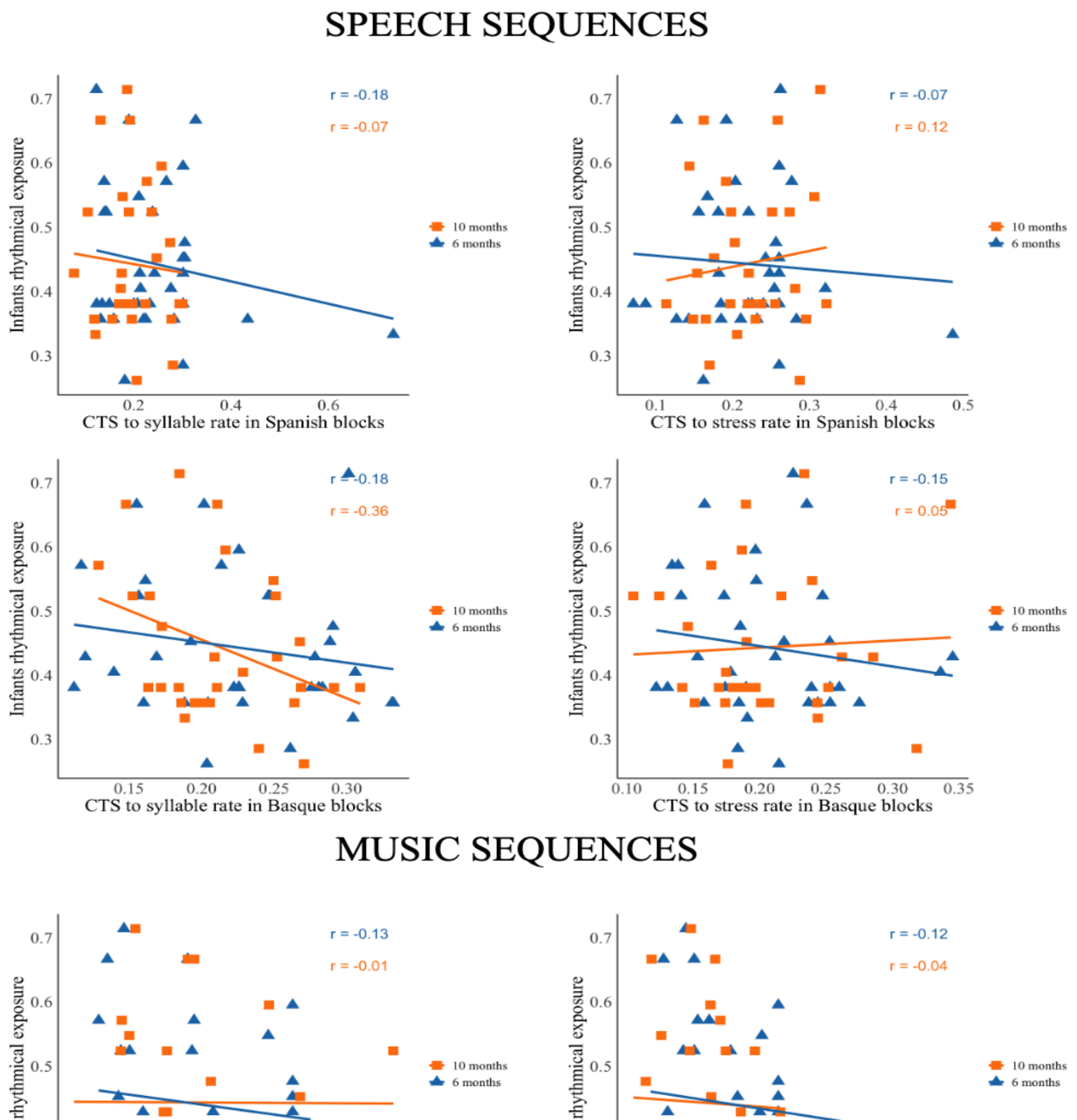

*Figure S1: Relationship between cortical tracking to speech and musical sequences in the stressed and syllable rates and infants' rhythmical exposure.*

### 2. ***Output of linear mixed effects models.***

*Table S1: Output of the linear mixed effects model on cortical tracking to musical sequences in the Spanish blocks in the delta band.*

| **Fixed effects** | ***ß*** | ***SE*** | ***df*** | ***t*** | ***p*** |
|---|---|---|---|---|---|
| Intercept | 1.0946 | 0.1319 | 126 | 8.300 | <.001 |
| Condition [Regular] | -0.1708 | 0.1762 | 94 | -0.969 | 0.335 |
| Age [Ten] | -0.1455 | 0.184 | 99 | -0.79 | 0.431 |
| Condition [Regular]:Age [Ten] | -365 | 0.259 | 94 | -1.41 | 0.162 |

*Note.* Number of observations = 134, ID (participant) = 37. Model = Coherence ~ Condition * Age + (1 |ID)

*Table S2: Output of the linear mixed effects model on cortical tracking to musical sequences in the Spanish blocks in the theta band.*

| **Fixed effects** | ***ß*** | ***SE*** | ***df*** | ***t*** | ***p*** |
|---|---|---|---|---|---|
| Intercept | 0.774 | 0.162 | 128 | 4.777 | <.001 |
| Condition [Regular] | 0.867 | 0.217 | 97 | 3.994 | <.001 |
| Age [Ten] | -0.272 | 0.228 | 102 | -1.194 | 0.235 |
| Condition [Regular]:Age [Ten] | -0.098 | 0.32 | 97 | -0.308 | 0.758 |

*Note.* Number of observations = 135, ID (participant) = 37. Model = Coherence ~ Condition * Age + (1 |ID)

*Table S3: Output of the linear mixed effects model on cortical tracking to musical sequences in the Basque blocks in the delta band.*

| **Fixed effects** | ***ß*** | ***SE*** | ***df*** | ***t*** | ***p*** |
|---|---|---|---|---|---|
| Intercept | 0.726 | 0.108 | 132 | 6.691 | <.001 |
| Condition [Regular] | 0.381 | 0.152 | 96 | 2.506 | 0.013 |
| Age [Ten] | -0.215 | 0.159 | 101 | -1.352 | 0.179 |
| Condition [Regular]:Age [Ten] | -0.362 | 0.225 | 96 | -1.609 | 0.11 |

*Note.* Number of observations = 136, ID (participant) = 37. Model = Coherence ~ Condition * Age + (1 |ID)

*Table S4: Output of the linear mixed effects model on cortical tracking to musical sequences in the Basque blocks in the theta band.*

| **Fixed effects** | ***β*** | ***SE*** | ***df*** | ***t*** | ***p*** |
|---|---|---|---|---|---|
| Intercept | 1.336 | 0.145 | 116 | 9.202 | <.001 |
| Condition [Regular] | 0.429 | 0.181 | 96 | 2.363 | 0.02 |
| Age [Ten] | -0.652 | 0.191 | 100 | -3.404 | <.001 |
| Condition [Regular]:Age [Ten] | 0.521 | 0.268 | 96 | 1.938 | 0.55 |

*Note.* Number of observations =136, ID (participant) = 37. Model = Coherence ~ Condition * Age + (1 |ID)

*Table S5: Output of the linear mixed effects model on cortical tracking to speech sequences in the Spanish blocks in the delta band.*

| **Fixed effects** | ***β*** | ***SE*** | ***df*** | ***t*** | ***p*** |
|---|---|---|---|---|---|
| Intercept | 1.615 | 0.1418 | 131 | 11.395 | <.001 |
| Condition [Regular] | -0.207 | 0.195 | 95 | -1.063 | 0.29 |
| Age [Ten] | -0.707 | 0.204 | 100 | -3.451 | <.001 |
| Condition [Regular]:Age [Ten] | 0.817 | 0.289 | 95 | 2.827 | <.001 |

*Note.* Number of observations = 136, ID (participant) = 37. Model = Coherence ~ Condition * Age + (1 |ID)

*Table S6: Output of the linear mixed effects model on cortical tracking to speech sequences in the Spanish blocks in the theta band.*

| **Fixed effects** | ***β*** | ***SE*** | ***df*** | ***t*** | ***p*** |
|---|---|---|---|---|---|
| Intercept | 1.187 | 0.146 | 131 | 8.116 | <.001 |
| Condition [Regular] | 0.438 | 0.205 | 96 | 2.133 | 0.035 |
| Age [Ten] | -0.13 | 0.214 | 100 | -0.611 | 0.542 |
| Condition [Regular]:Age [Ten] | -0.573 | 0.303 | 96 | -1.892 | 0.06 |

*Note.* Number of observations = 135, ID (participant) = 37. Model = Coherence ~ Condition * Age + (1 |ID)

*Table S7: Output of the linear mixed effects model on cortical tracking to speech sequences in the Basque blocks in the delta band.*

| **Fixed effects** | ***β*** | ***SE*** | ***df*** | ***t*** | ***p*** |
|---|---|---|---|---|---|

| | | | | | |
|---|---|---|---|---|---|
| Intercept | 1.168 | 0.108 | 129 | 10.791 | <.001 |
| Condition [Regular] | 0.137 | 0.146 | 97 | 0.941 | 0.349 |
| Age [Ten] | 0.096 | 0.153 | 101 | 0.626 | 0.532 |
| Condition [Regular]:Age [Ten] | -0.395 | 0.216 | 97 | -1.829 | 0.07 |

*Note.* Number of observations = 136, ID (participant) = 37. Model = Coherence ~ Condition * Age + (1 |ID)

*Table S8: Output of the linear mixed effects model on cortical tracking to speech sequences in the Basque blocks in the theta band.*

| **Fixed effects** | ***β*** | ***SE*** | ***df*** | ***t*** | ***p*** |
|---|---|---|---|---|---|
| Intercept | 1.055 | 0.117 | 123 | 8.973 | <.001 |
| Condition [Regular] | 0.413 | 0.151 | 95 | 2.72 | .007 |
| Age [Ten] | 0.054 | 0.160 | 99 | 0.340 | .734 |
| Condition [Regular]:Age [Ten] | -0.37 | 0.224 | 95 | -1.655 | .101 |

*Note.* Number of observations =135, ID (participant) = 37. Model = Coherence ~ Condition * Age + (1 |ID)

#### 3. *Spectral features of the stimuli.*

*Figure S2: Acoustic properties of Spanish and Basque musical and speech sequences. Here, we show the spectral information of speech and musical sequences in normalized power (scaled power of signal). Spectra on the left correspond to Spanish stimuli, showing peaks at 2 and 4 Hz for Regular condition. Spectra on the right correspond to Basque stimuli, showing peaks at 1.6 and 4 Hz for Regular condition.*

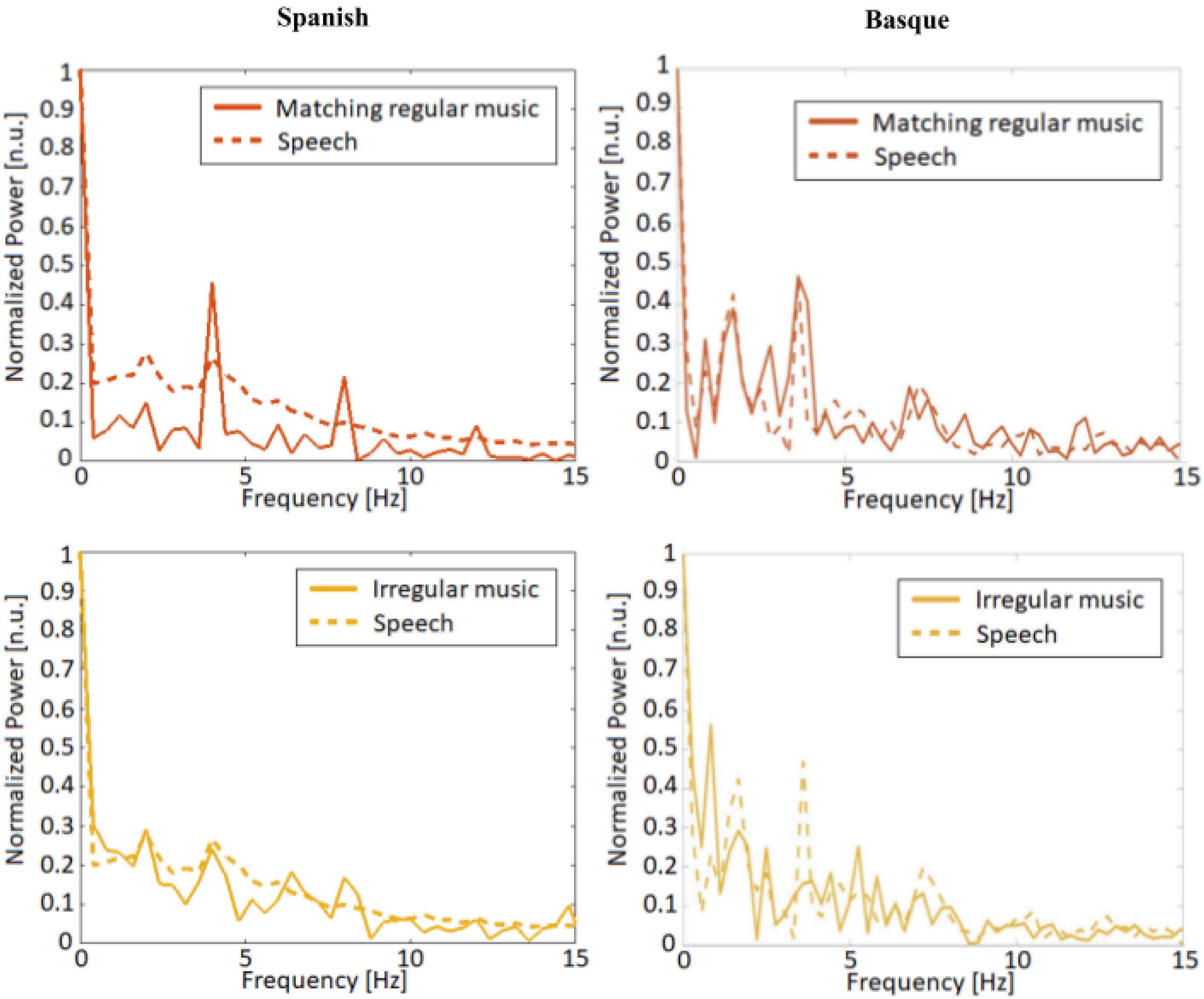

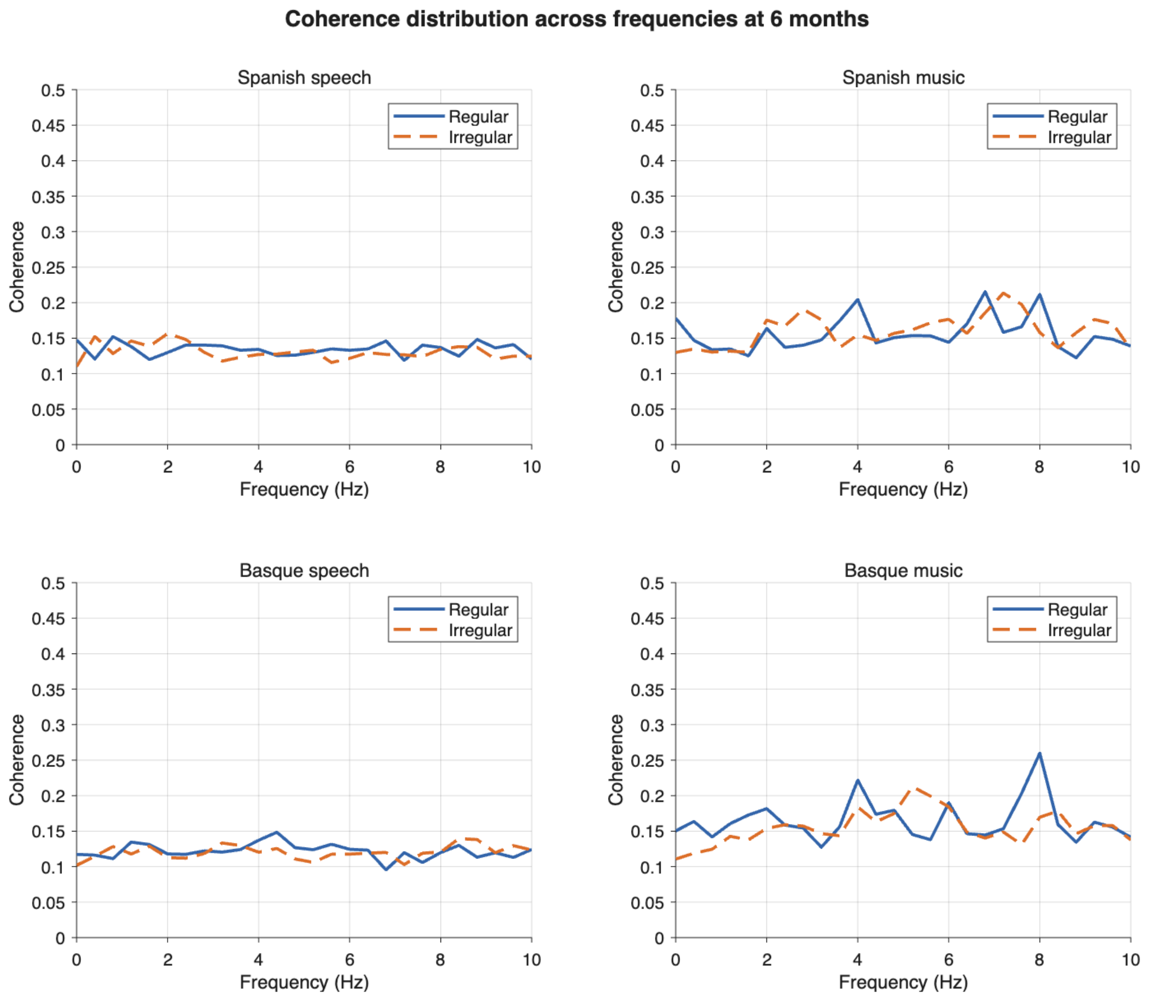


*Figure S3: Coherence between EEG and auditory stimuli in 6-month-old infants. Here, we show the coherence spectra averaged across all electrodes and participants for Spanish and Basque speech and music sequences. Coherence values reflect the degree of neural synchronization between the infant EEG signal and the auditory envelope of each stimulus. Spectra on the top correspond to Spanish stimuli, for both speech and music conditions. Spectra on bottom correspond to Basque stimuli, for both speech and music conditions. In each panel, the Regular condition (solid line) and Irregular condition (dashed line) are shown across frequencies from 0 to 10 Hz.*

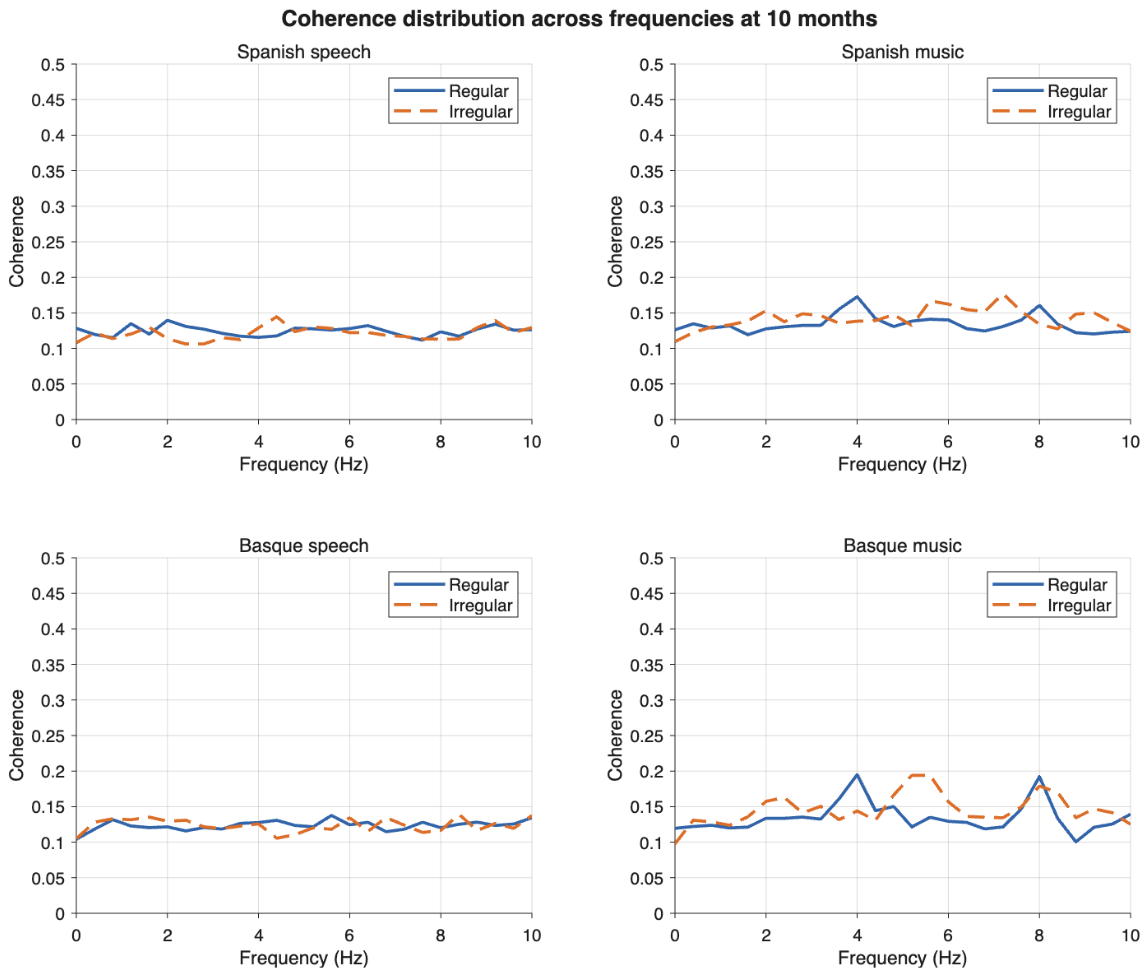


*Figure S4: Coherence between EEG and auditory stimuli in 10-month-old infants. Here, we show the coherence spectra averaged across all electrodes and participants for Spanish and Basque speech and music sequences. Coherence values reflect the degree of neural synchronization between the infant EEG signal and the auditory envelope of each stimulus. Spectra on the top correspond to Spanish stimuli, for both speech and music conditions. Spectra on bottom correspond to Basque stimuli, for both speech and music conditions. In each panel, the Regular condition (solid line) and Irregular condition (dashed line) are shown across frequencies from 0 to 10 Hz.*
